# Host PDZ-containing proteins targeted by SARS-Cov-2

**DOI:** 10.1101/2021.02.01.429176

**Authors:** Célia Caillet-Saguy, Fabien Durbesson, Veronica V. Rezelj, Gergö Gogl, Quang Dinh Tran, Jean-Claude Twizere, Marco Vignuzzi, Renaud Vincentelli, Nicolas Wolff

## Abstract

Small linear motif targeting protein interacting domains called PDZ have been identified at the C-terminus of the severe acute respiratory syndrome coronavirus 2 (SARS-CoV-2) proteins E, 3a, and N. Using a high-throughput approach of affinity-profiling against the full human PDZome, we identified sixteen human PDZ binders of SARS-CoV-2 proteins E, 3A and N showing significant interactions with dissociation constants values ranging from 3 μM to 82 μM. Six of them (TJP1, PTPN13, HTRA1, PARD3, MLLT4, LNX2) are also recognized by SARS-CoV while three (NHERF1, MAST2, RADIL) are specific to SARS-CoV-2 E protein. Most of these SARS-CoV-2 protein partners are involved in cellular junctions/polarity and could be also linked to evasion mechanisms of the immune responses during viral infection. Seven of the PDZ-containing proteins among binders of the SARS-CoV-2 proteins E, 3a or N affect significantly viral replication under knock-down gene expression in infected cells. This PDZ profiling identifying human proteins potentially targeted by SARS-CoV-2 can help to understand the multifactorial severity of COVID19 and to conceive effective anti-coronaviral agents for therapeutic purposes.

## Introduction

Coronaviruses (CoV) are enveloped viruses that primarily infect birds and mammals. Some CoVs have evolved to establish zoonotic disease in humans, potentially causing severe respiratory illnesses [1]. The seriousness of CoV infection during the last outbreaks caused by the severe acute respiratory syndrome coronavirus 1 (SARS-CoV) in 2003; the Middle East Respiratory Syndrome-virus (MERS-CoV) in 2012 and the SARS-CoV-2 in 2020 [2], and the lack of effective treatment for CoV infection highlight the need of a deeper understanding of coronavirus molecular biology to elaborate therapeutic strategies that counteract SARS-CoV-2 infection. However, limited knowledge of the molecular details of SARS-CoV-2 infection precludes a fast development of compounds targeting the host-virus interface. Several SARS-CoV-2 interactomes have been recently published using mass spectrometry [3–5]. Unfortunately, some transient and weak interactions with affinities in the 1-100 μM range can be hardly detected by such approaches. However, during infection, viruses target host signaling pathways that mainly involve protein-protein interactions of low affinity, as transient bindings are essential for the dynamics of cellular homeostasis.

Hijacking of cellular protein functions is a widely used strategy for viruses as they are obligate intracellular pathogens. Each step of the viral life cycle from entry to transmission is orchestrated through interactions with cellular proteins. All families of viruses manipulate the cell proteome by targeting key proteins involved in the control of cell defense and homeostasis. Interactions mediated by short linear motifs (SLiMs) are ubiquitous in the eukaryotic proteome [6] and the adaptation of viruses to their environment involves the extensive use of SLiM mimicry to subvert host functionality [7].

PDZ-Binding Motifs (PBMs) are SLiMs that interact with a large family of protein interaction domains called PDZ (PSD-95/Dlg/ZO-1). PBMs are usually located at the extreme carboxyl terminus of proteins [8]. The PDZ-PBM interactome is one of the most prominent instances of a SLiM-mediated protein interaction network serving key cell signaling purposes [9]. PBMs were identified in diverse proteins of many viruses responsible for acute to chronic infections [10,11]. Cellular PDZ-viral PBM interactions were shown to be directly involved in viral pathogenicity of SARS-CoV, as well as of rabies virus and in the oncogenicity of human papillomavirus (HPV16) [10]. During infection, the viral proteins compete with the endogenous ligands through the binding to the PDZ domain of the host protein targets allowing a cellular relocalization, sequestration or degradation of protein [12] but can also affect the catalytic activity of signaling proteins [13]. Functional perturbations of cellular processes due to interactions between viral PBMs and cellular PDZ-containing proteins can facilitate the virus life cycle in the host and the dissemination to new hosts. Thus, targeting the PBM-PDZ interface could potentially lead to novel antiviral therapies.

The four major structural proteins encoded by the coronavirus genome; the spike (S) protein, the nucleocapsid (N) protein, the membrane (M) protein, and the envelope (E) protein, participate in different aspects of the replication cycle, and ultimately form a structurally complete viral particle. While a minor fraction is incorporated into the virion envelope, the integral membrane E protein (~8–12 kDa) of CoVs including SARS-CoVs is a multifunctional protein that forms ion channels with its transmembrane (TM) domain. This channel activity is mediated by formation of pentameric oligomers and is only mildly selective for cations [14]. The channel would be active in the secretory pathway, altering luminal environments, rearranging secretory organelles and leading to efficient trafficking of virions [15]. In addition, E protein interacts with host proteins, related to its extramembrane domain particularly the C-terminal domain.

The C-terminal sequence in extra membrane position is predicted to be partially folded (helices and ß-coil-ß motif) which could affect the global conformation of the pentameric E channel suggesting that the channel activity of E and its protein interactions most likely interplay [14,16]. In addition, the E protein encodes a putative PBM sequence at its extreme C-terminus (Fig. 1A) that could also interfere through protein-protein interactions with the oligomeric organization of E protein and consequently its channel activity. The PBM sequence of E protein has been identified as a virulence factor for SARS-CoV virus, influencing replication level, virus dissemination, and pathogenicity of SARS-CoV in animal models [17]. To date, E protein of SARS-CoV has only been reported to interact with five host proteins; Bcl-xL, PALS1/MPP5, syntenin, sodium/potassium ATPase a1 subunit and stomatin [16]. The relevance of these interactions is not yet fully understood. Among these five partners of SARS-CoV, PDZ domains of the PALS1/MPP5 tight junction protein and of Syntenin interacts with the C-terminal PBM of E protein. PBM-PDZ interactions strongly impact the structure of mammalian epithelial cells and could contribute to the abrupt deterioration of the lung epithelium observed in patients infected by SARS-CoV [18]. Therefore, to better understand the influence of host PDZ and viral PBM motifs in the pathogenicity of SARS-CoV-2, we performed a targeted screen to identify cellular PDZ domains capable of interacting with PBM motifs CoV proteins.

**Figure 1:**
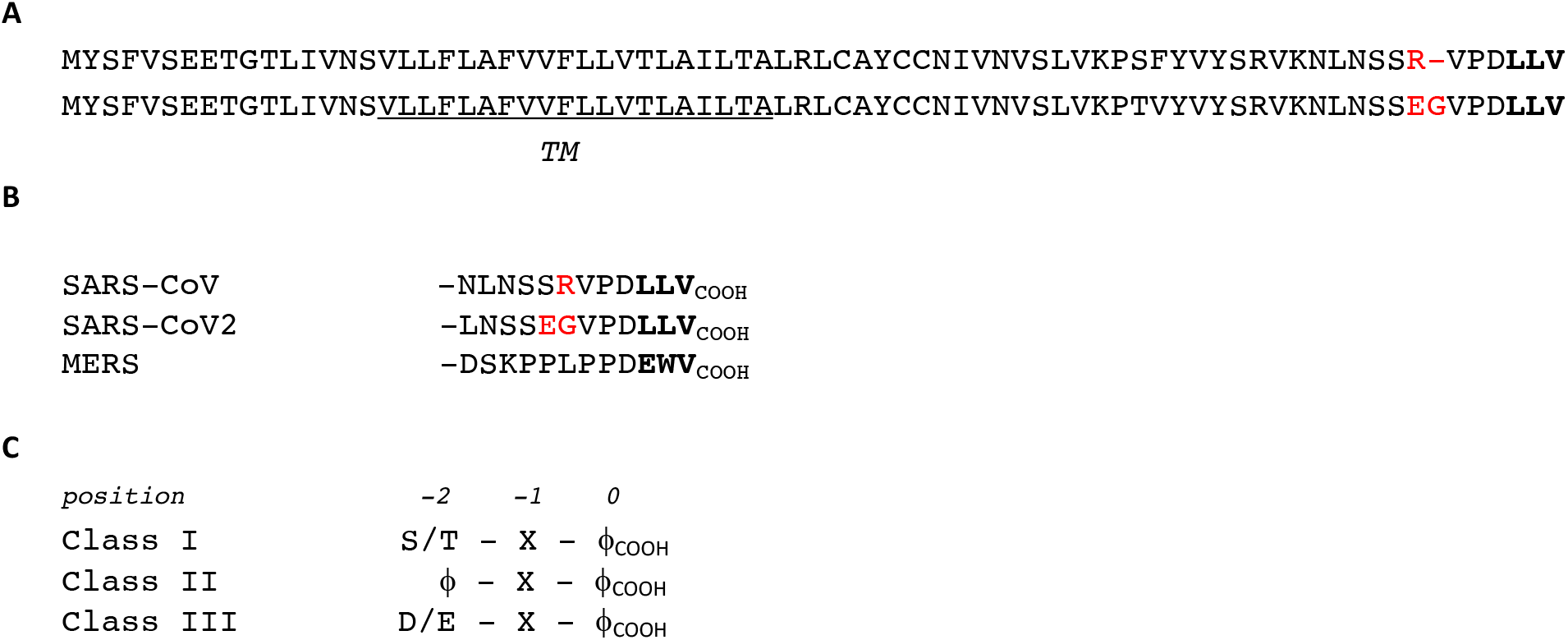
E protein sequences. (A) Comparison of E protein sequences of SARS-CoV-2 and SARS-CoV viruses. The SARS-CoV-2 genome is very similar to SARS-CoV. The last three residues that form the PBM sequence (in bold) are strictly conserved. Deletion and substitution of amino-acid close to the PBMs are colored in red. The transmembrane helix (TM) is underlined. (B) Sequences of peptides used as baits in the holdup assays. PBM residues are in bold. (C) The three classes of PBM. X is any amino-acid (position - 1), and φ is a hydrophobic amino-acid (position 0).

In this study, we used the holdup assay [19], a high-throughput and quantitative approach that offers a high sensitivity for low-to-medium (1-100 μM) affinity, typically found for PDZ/PBM interactions. We established the list of PDZ-containing partners potentially targeted by SARS-CoV-2 E protein through its PBM. This work identified ten PDZ and determined their affinities to SARS-CoV-2 E protein. Most of them are involved in cellular junction dynamics (PARD3, TJP1/ZO-1, LNX2, MLLT4, NHERF1). To further understand the potential implication of the PDZ-PBM in pathogenicity, we have also determined the PDZ-containing proteins recognized by the E protein of SARS-CoV and MERS-CoV viruses, as well as the PDZ binders of the SARS-CoV-2 3a and N proteins that also encode for a C-terminal PBM sequence. From this PDZ list, we have selected 20 PDZ-containing partners of SARS-CoV-2 to assess their influence on SARS-CoV-2 replication in infected cells and found that the expression of seven of them are relevant for the efficiency of viral replication.

## RESULTS

### SARS-CoV2 E protein recognizes several cellular PDZ-containing proteins

The E protein of SARS-CoV virus is known to contain a C-terminal motif, a - LLV_COOH_ sequence corresponding to a PBM of class II (Fig. 1C). This motif is strictly conserved in SARS-CoV2 E protein (Fig. 1A). To quantify the binding activity of the SARS-CoV2 E PBM, we used an *in vitro* automated high-throughput chromatography assay called holdup [19,20]. A 12-mer peptide was synthetized encompassing the C-terminal PBM sequence of SARS-CoV2 E protein linked to a biotinyl group (Fig. 1B). This peptide was used as bait to quantify the interaction between the SARS-CoV2 E protein PBM peptide and the full human PDZ domain library expressed in *Escherichia coli*. The updated PDZ library called PDZome V.2 covers 98 % of the human “PDZome” (266 over 272 PDZ domains identified in the human genome) [20,21]. Each PDZ of the library is defined by the protein name followed by the PDZ number in the protein. The holdup approach displays a high sensitivity of detection for low–to-medium affinity PDZ/PBM pairs and provides an affinity-based ranking of the identified binders corresponding to a specificity profile [22].

The specificity profiles for SARS-CoV, SARS-CoV-2 and MERS-CoV E PBM with the mean values of binding intensities (BI) are reported in Fig. 2A with an affinity-based ranking. Steady state holdup binding intensities can be converted to dissociation constants as described previously [23]. For this, we determined the dissociation constants of a few PBM-PDZ interactions using a competitive fluorescence polarization assay. The holdup assay has detected several significant interactions for the three viruses. Using a stringent threshold previously defined for significant detectable binders in holdup assays at BI > 0.2 (~100 μM dissociation constant), ten PDZ binders of SARS-CoV-2 E protein showed interactions with affinities values ranging from 2.6 μM (TJP1_2) to 82 μM (MLLT4) (Fig. 2, zoomed-in view, Table 1). Thus, the PBM sequence of SARS-CoV-2 E protein has the capacity to bind *in vitro* several PDZ domains in the 1-100 μM affinity range typically found for PDZ/PBM interactions. As illustrated by the sharp profile of interaction of Fig. 2A, SARS-CoV2 E PBM recognizes only 4 % of PDZ domains from the human PDZ library reflecting the specificity of these interactions.

**Figure 2;.**
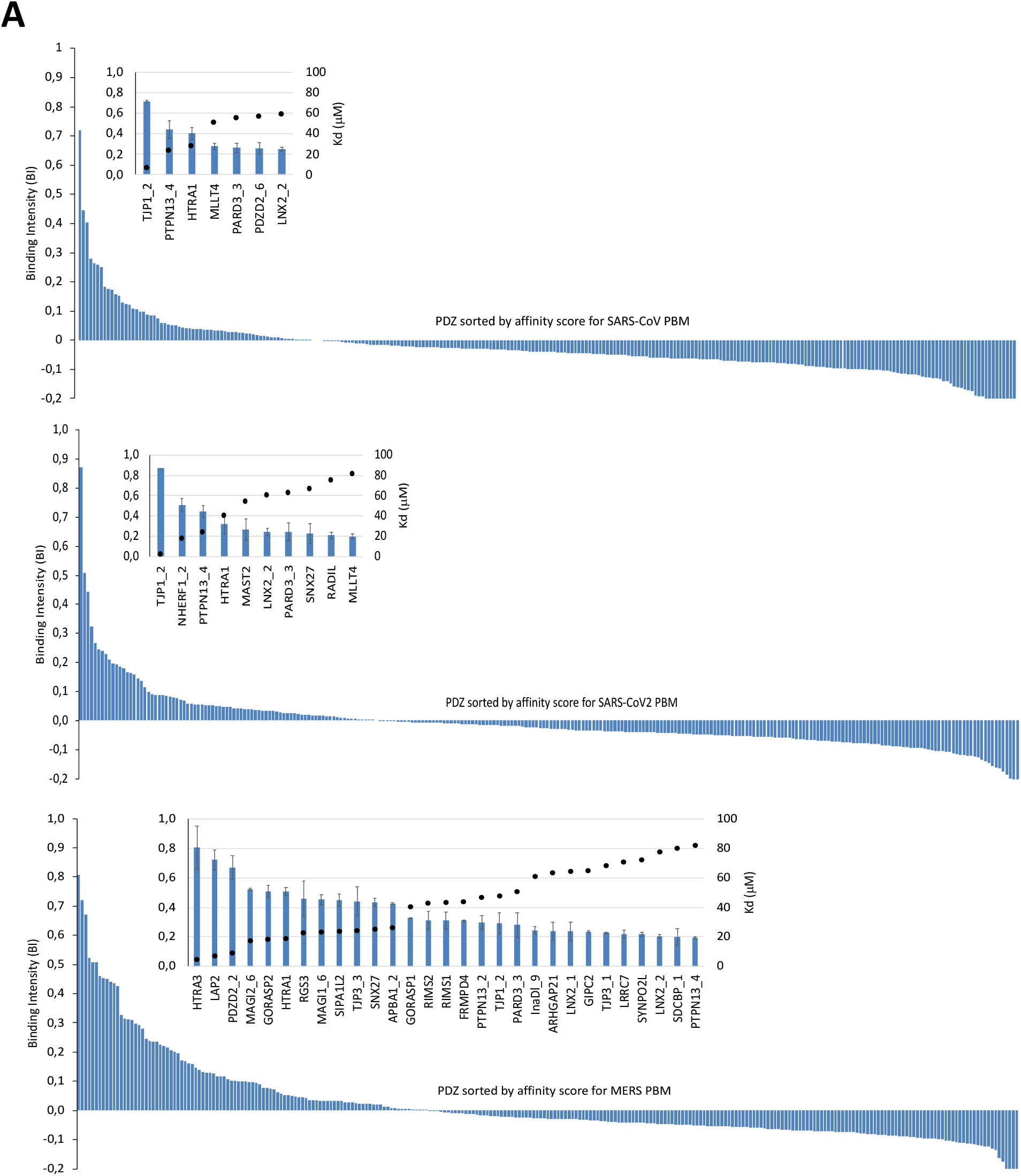
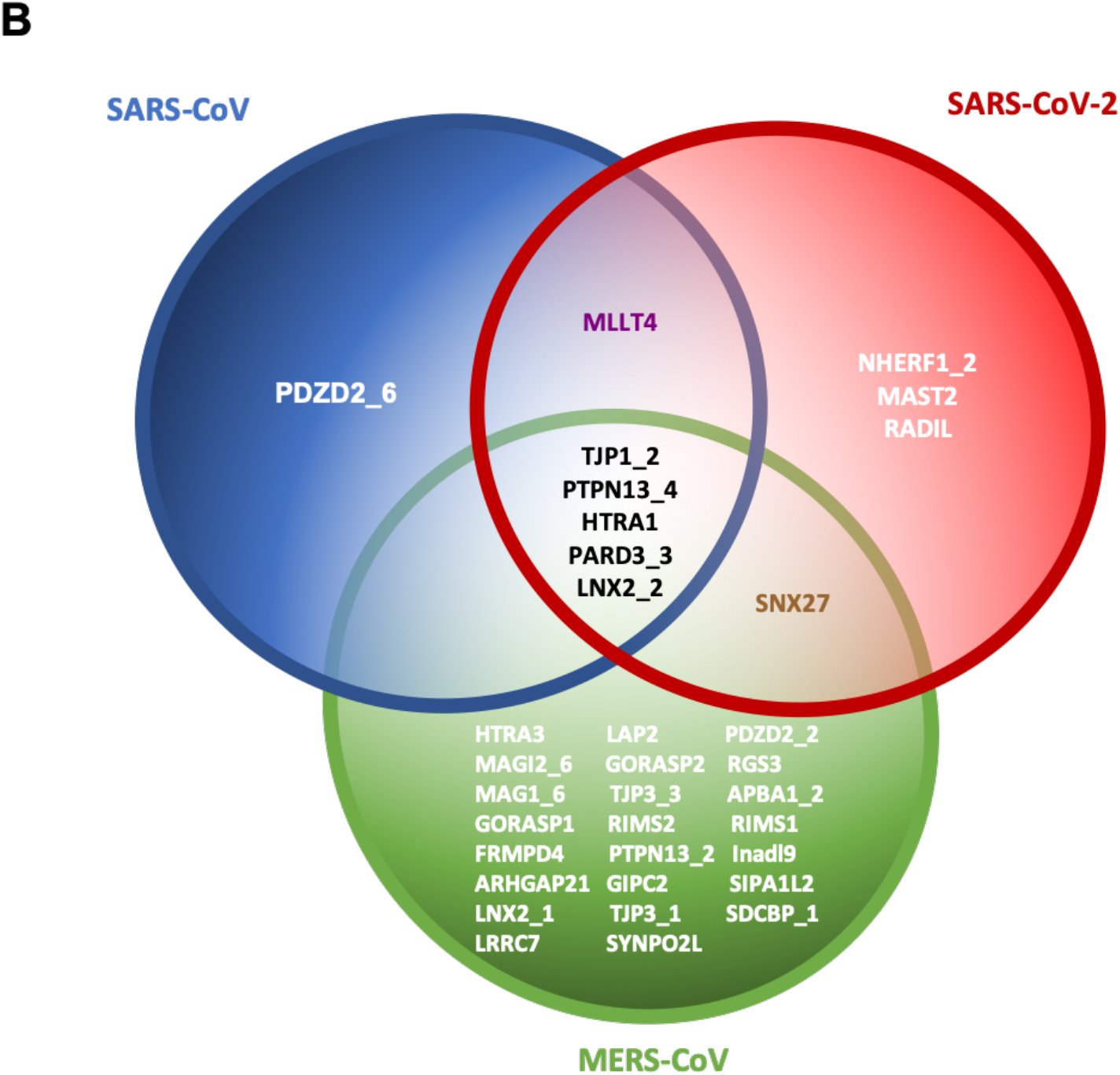
PDZ binders of the E protein PBMs by holdup assay. (A) PDZome binding profiles of the E protein PBMs by holdup assay of SARS-CoV (upper panel), SARS-CoV-2 (middle panel) and MERS-CoV (bottom panel). PDZ domains are ranked by decreased BI values. Zoomed-in views show the 7, 10 and 29 PDZ binders of SARS-CoV, SARS-CoV-2 and MERS-CoV E proteins, respectively, which displayed significant BI values higher than 0.2. Error bars are standard deviations of two independent experiments. The black circles correspond to affinity values (Kd) in μM (secondary Y axis). (B) Venn diagram of PDZ binders of the E protein PBMs of SARS-CoV, SARS-CoV-2 and MERS-CoV.

**Table 1;.**
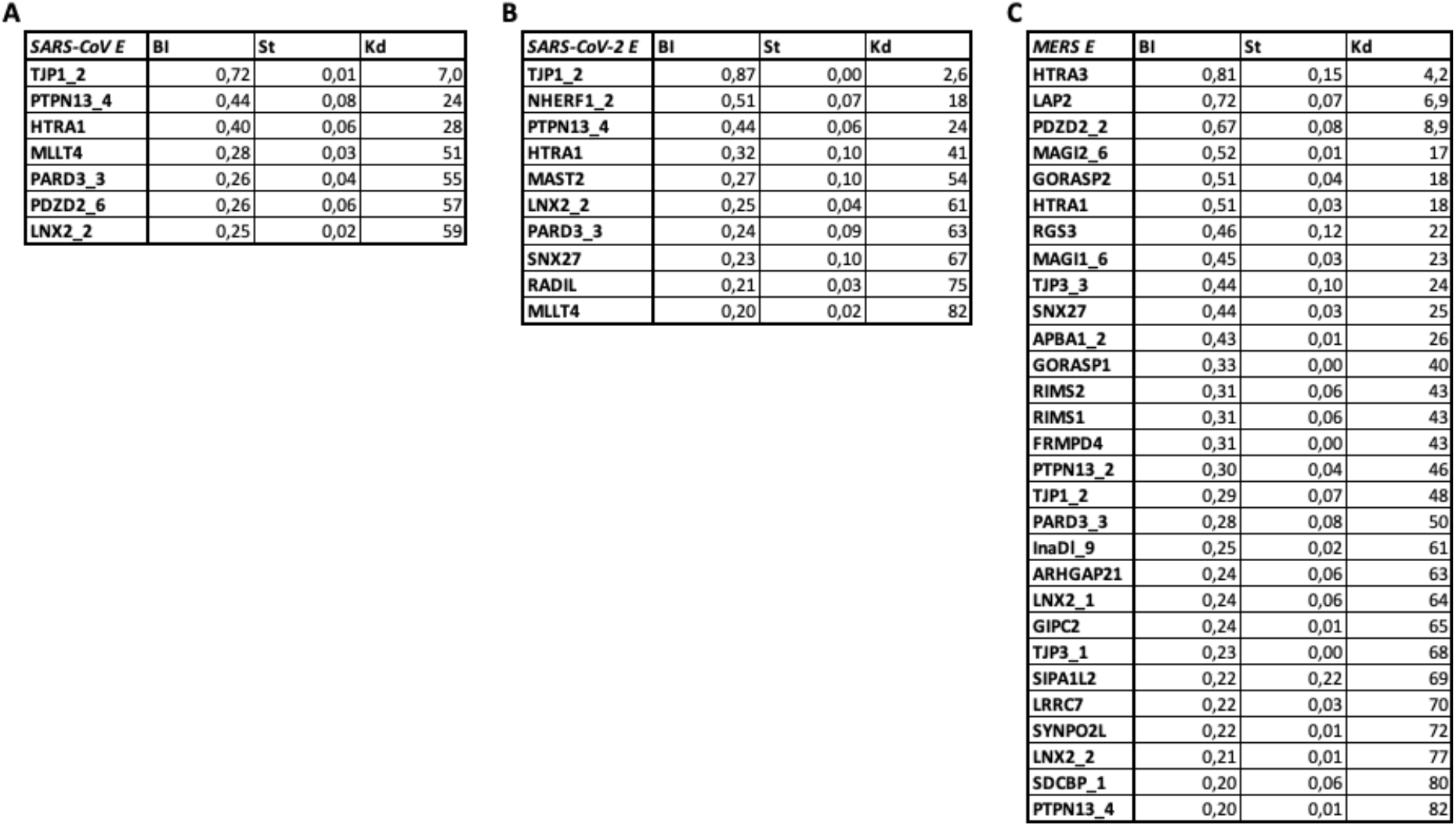
Measured BI and converted Kd (μM) values of PDZ domains for the SARS-CoV (A), SARS-CoV-2 (B) and MERS-CoV (C) E protein PBMs. Only significant BI values higher than 0.2 are reported. When multiple holdup experiments were performed, the averaged BI values and standard deviations are reported.

### Patterns of human PDZ-containing proteins preferentially targeted by E proteins expressed by SARS-CoV, SARS-CoV-2 and MERS-CoV viruses

Two differences of sequence between SARS-CoV and SARS-CoV2 E proteins are observed upstream their identical PBM sequences (Fig. 1B).

Seven PDZ-containing proteins (3 %) were identified as binders of SARS-CoV E PBM by high-throughput holdup (Table 1). Six of them are also SARS-CoV-2 E PBM binders, ranked by decreased affinity values in Fig. 2B; TJP1-2, PTPN13_4, HTRA1, MLLT4, PARD3_3 LNX2_2. The affinities of SARS-CoV and SARS-CoV-2 E PBMs for this set of proteins are similar, in agreement with the strict conservation of the last five C-terminal positions of the PBM (p0 to p-4)(Fig. 1A; Table 1). PDZ2_6 is only recognized by SARS-CoV E PBM while four additional PDZs are only recognized by SARS-CoV-2 E PBM (NHERF1_2, MAST2, RADIL, SNX27). Thus, the PBM upstream sequence differences increase the number of binders for SARS-CoV-2 protein E with four additional host partners.

The MERS-CoV E protein encodes for a class III PBM (Fig. 1B and C). As compared to the SARS-CoV/ SARS-CoV-2 E patterns, the profile of interaction of MERS-CoV E PBM is extended with a larger number of partners, i.e. 29 (11 %), within the same range of affinities between 5 μM and 82 μM (Table 1). Six over seven PDZ-containing partners of SARS-CoV E PBM, and six over ten PDZ-containing partners of SARS-CoV-2 E PBM are also recognized by MERS-CoV E PBM (Fig. 2B).

### The C-terminal motifs of SARS-CoV2 N and 3a proteins also bind to cellular PDZ-containing proteins

We have systematically analyzed the sequences of the 29 proteins of SARS-CoV-2 to search for potential PBM. Consistent with previous work on SARS-CoV [24], we identified two PBM motifs at the C-terminus of two SARS-CoV-2, namely proteins E and 3a. The PBM sequences of E and 3a proteins (Fig. 3A), two viroporins, belong to the same class of PBM, the class II (Fig. 1C). While the PBM of the E protein is involved in SARS-CoV virulence and replication [17], the motif of the protein 3a has no significant impact on viral growth [24]. However, it has been shown that this accessory 3a protein might exhibit compensatory function in the absence of E, on viral replication *in vitro* and *in vivo*. This homology suggests that the E and 3a proteins could interact with common host PDZ-containing proteins.

**Figure 3;.**
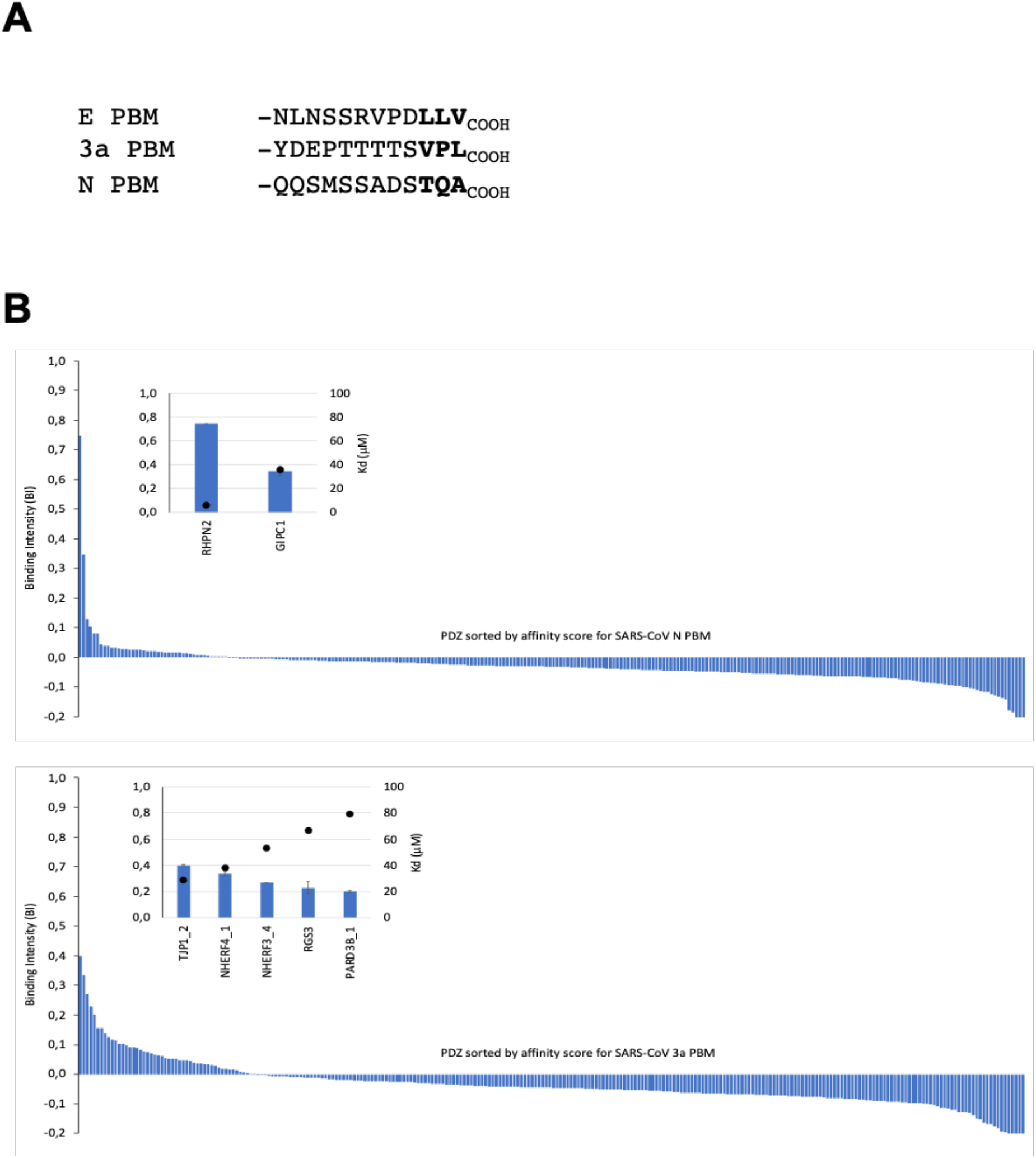
PDZome binding profiles of SARS-CoV2 N protein and 3a protein PBM by holdup assay. (A) sequences of E, N and 3a peptides used as baits in the holdup assays. (B) PDZ domains are ranked by decreased BI values. Zoomed-in views show the 2 and 5 PDZ binders of SARS-CoV-2 N and 3a proteins, respectively, which displayed significant BI values higher than 0.2. The black circles correspond to affinity values (Kd) in μM (secondary Y axis).

We also identified a potential PBM sequence at the C-terminus of the structural nucleocapsid protein N. This class I motif is atypical by the presence of a small hydrophobic residue, i.e an alanine, at the last position (Fig. 3A).

The binding profiles of PBM from SARS-CoV-2 3a and N proteins are reported in Fig. 3B. Five binders (2%) displayed significant BI values higher than 0.2: TJP1_2, NHERF4_1, NHERF3_4, RGS3 and PARD3B_1, ranked by decreased affinity values. Interestingly, E protein of SARS-CoV and SARS-CoV-2 and 3a protein of SARS-CoV-2 have the same best binder TJP1_2, and both also recognized the PDZ domain of PARD3B. NHERF3 and NHERF4 are two new binders of SARS-CoV-2 while RGS3 is also recognized by MERS-CoV E protein. To note, the affinity range of 3a protein for its binders is lower than the one of E protein, from 25 to 80 μM. Concerning the N protein PBM binders, only two PDZ domains (1 %) are identified: RHNP2 and GIPC1. These proteins have not been identified as binders of proteins E and protein 3a. GIPC1 is a PDZ-containing protein recognized by the PBM of the hepatitis B virus capsid protein and can accommodate another atypical small C-terminal residue, a cysteine [25].

### Effects of depletion of cellular PDZ-containing proteins on SARS-CoV2 replication

We next investigated the biological role of the PDZ-containing partners of SARS-CoV-2 proteins in the viral replication according to our holdup assay results.

We established a list of potential PDZ partners recognized by at least one of the PBM motif encoded by the SARS-CoV-2 proteins E, 3a or N (Table 2). We selected the 16 PDZ partners identified with binding intensities higher than 0.2 (Fig. 2B and 3B), and added 4 additional PDZ partners with BI values comprised between 0.16 and 0.20; MPP5/PALS1, PDZD2 and DLG3 recognized by SARS-CoV-2 E protein and FRMPD4, a binder of both SARS-CoV-2 proteins E and 3A.

**Table 2;.**
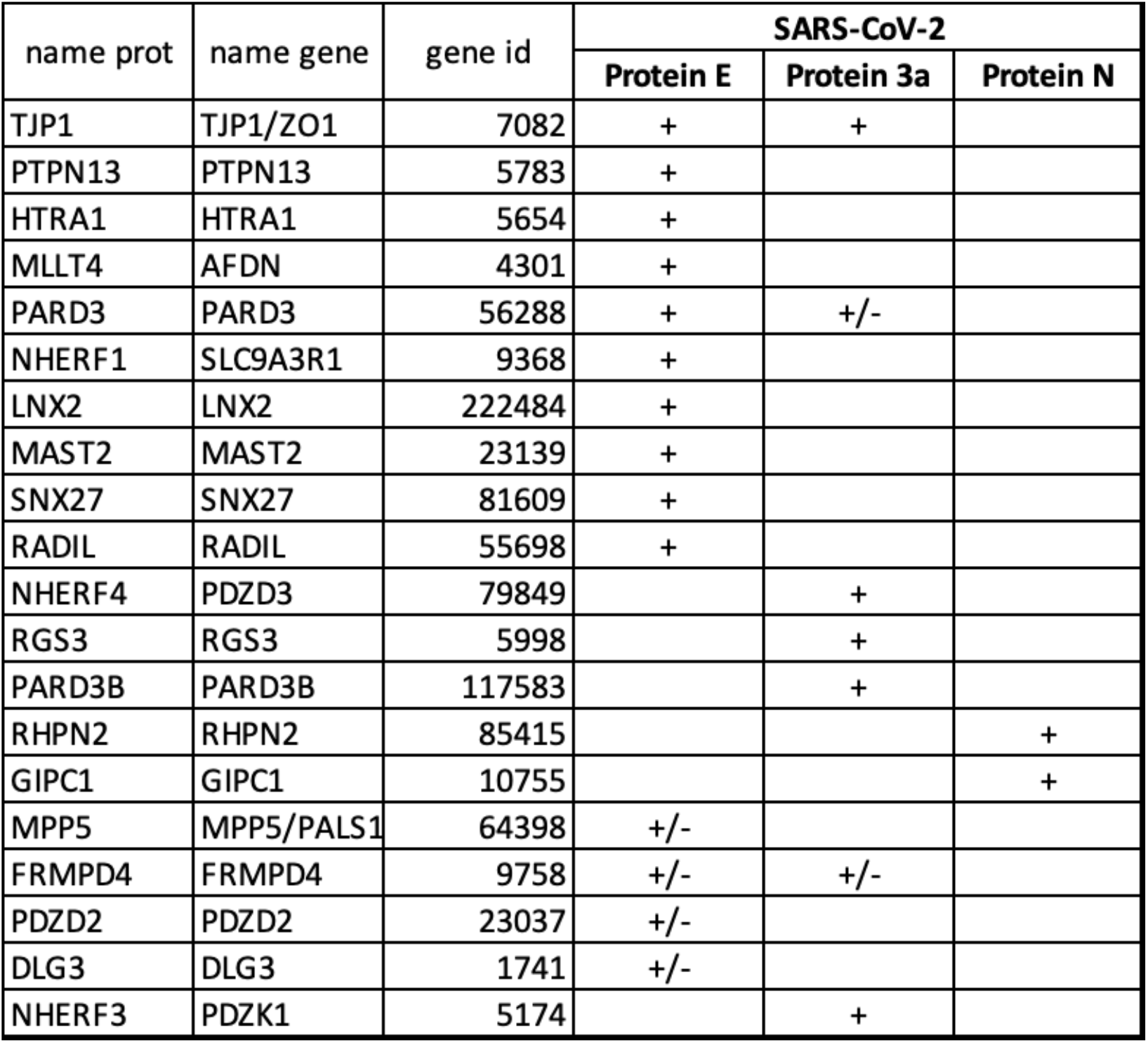
Cellular PDZ-containing proteins recognized by at least one SARS-CoV-2 protein in holdup assays. List of the 20 PDZ partners of SARS-CoV-2 proteins selected for knock-down by siRNA transfection in human lung A549 cells. (+) indicated PDZ partners identified by assay with binding affinity values higher than 0.2; (+/-) represented PDZ binders with BI values comprised between 0.16 (105 μM) and 0.20 (80 μM).

Twenty human PDZ-containing proteins were targeted for knock-down by siRNA transfection in human lung A549 cells. This cell line stably expresses the SARS-CoV-2 receptor ACE2 [26]. Our approach to knock-down gene expression used siRNA pools containing a mixture of four individual siRNAs), that are designed and modified for greater specificity, providing guaranteed gene silencing of human targets. We have validated these siRNA pools previously in a screen carried out in human A549 cells overexpressing ACE-2 (Table S1). Our approach confirmed that knockdown of a number of different genes, including the receptor ACE-2, and other unexpected genes such as BZW2 or BRD4 reduces SARS-CoV-2 replication in these cells [3]. The effect of PDZ partner knock-down on virus replication was assessed in three independent experiments and compared to virus replication in control siRNA-transfected cells (two-way ANOVA with Dunnet’s test). As expected, knock-down of the SARS-CoV-2 receptor ACE2 resulted in significantly lower virus replication (Fig. 4). Virus replication was also significantly reduced in PARD3 and RSG3 knocked-down cells, suggesting a pro-viral role for these human proteins. PARD3 is found in our holdup assays as a PDZ partner of E protein of SARS-CoV, SARS-CoV-2, and MERS-CoV while RGS3 is recognized by SARS-CoV2 3a protein. Comparatively, knock-down of MLLT4, PARD3B, MAST2, RHPN2 and HTRA generated an increased virus replication. MLLT4, MAST2, and HTRA1 interact with SARS-CoV-2 E protein, PARD3B with SARS-CoV-2 3a protein, and finally RHPN2 binds to SARS-CoV-2 N protein.

**Figure 4;.**
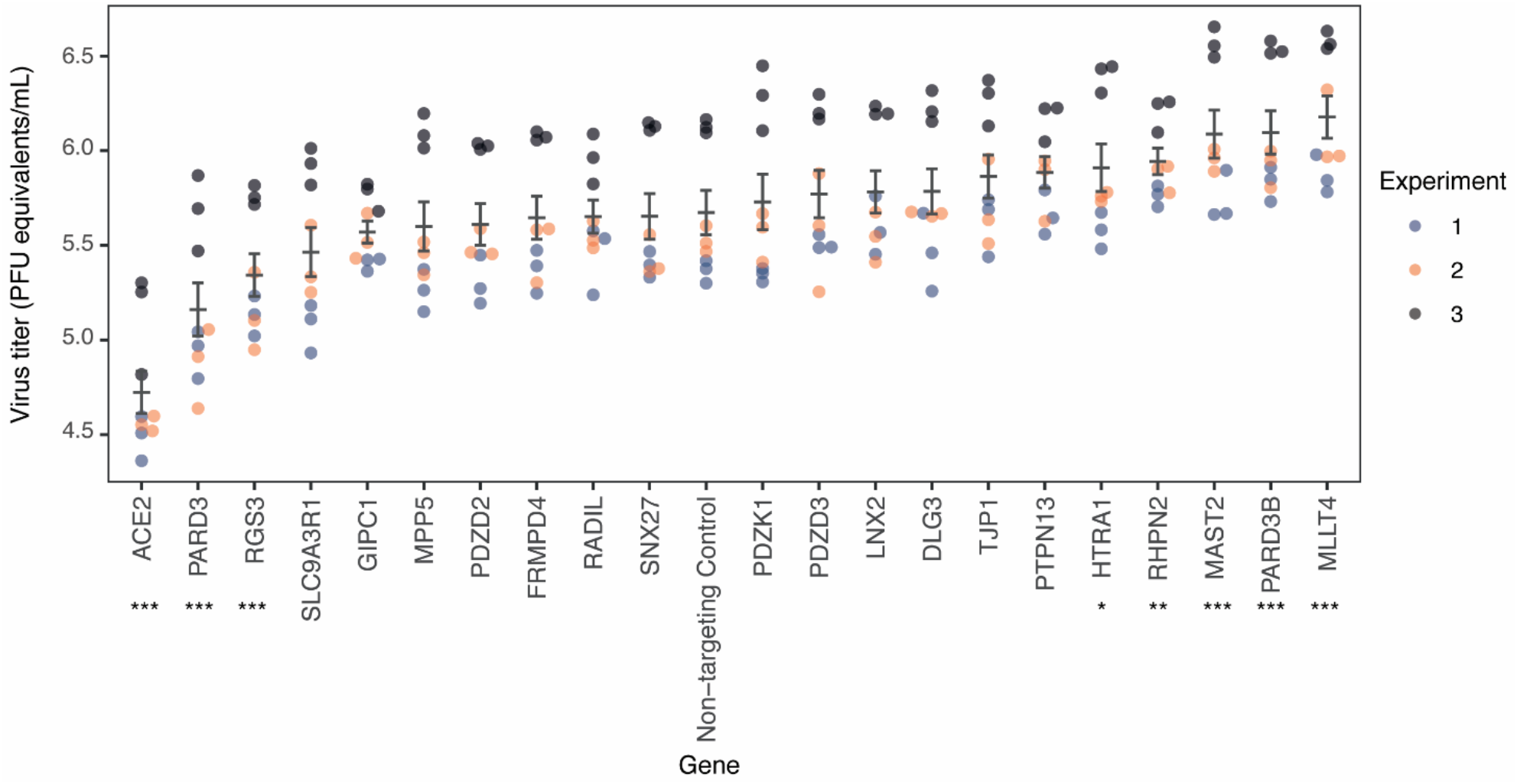
Effect of knock-down of human PDZ-containing proteins on SARS-CoV-2 replication. The indicated genes were knocked down by siRNA transfection in human lung A549-ACE2 cells. ACE2 knockdown was used as a positive control. Virus load in knocked-down cells was assessed in three independent experiments 72h post infection and is shown as plaque forming units (PFU) equivalents/mL supernatant. The virus load in each condition was compared to that in control siRNA-transfected cells (two-way ANOVA with Dunnett’s test for multiple comparisons \*\*\**p*≤0.001; \*\**p*=0.00611; \**p*=0.02672).

Altogether, our results suggested that the expression of these seven proteins is relevant for viral replication.

## DISCUSSION

Only a few partners of SARS-CoV-2 E protein have been identified through extensive high throughput approaches such as proteomics. As an example, among the structural SARS-CoV-2 proteins, only 6 binders of E protein are found as monitored in the Gordon and *al*. study, while 30 partners for M protein and 15 binders to N proteins are identified [3]. Among them, no PDZ-containing protein has been identified in this study. Here, we have focused on specific interactions involving short C-terminal motifs of SARS-CoV-2 proteins, the PBMs, and PDZ domains of human partners. We mainly used the holdup high-throughput screening to identify the host PDZ partners of E, 3a and N proteins and to quantify their PDZ-PBM affinities and specificities [19]. This chromatographic assay in solution is fully suitable to accurately rank low affinity interactions in the submillimolar range, which may escape detection by other high-throughput methods.

### Conservation of the PBM sequence in Coronavirus E protein

The PBM motif at the C-terminus of E protein is only expressed by coronoviral genera a and ß and is absent from the genus g. The last 4 residues of E protein are strictly conserved in the SARS-CoV and SARS-CoV-2 viruses. A recent study has shown that, among the 3617 available complete genome sequences of SARS-CoV-2 in the NCBI database, only two variants (DFLV and YLLV) have been reported in the DLLV PBM motif of the E protein [27].

Both the E proteins of SARS-CoV-2 and MERS-CoV comprised putative PBM sequences at their C-terminus but their nature is very different with a class II PBM for the genus a, and a class III PBM for the genus ß. Our work shows that despite the absence of conservation between the PBM sequence of SARS-CoV/CoV-2 and MERS-CoV E protein, the two motifs interact with a common set of host partners; indeed, among the ten host partners we identified for SARS-CoV-2, six are also binders of MERS-CoV.

### Host-viral protein-protein interactions

#### Comparison of host partners of CoV and CoV-2

The affinity-based ranking of the identified binders of E protein provided a specificity profile. As comparison, the class I PBMs of NS5 protein from West Nile virus and of capsid protein from hepatitis B virus interact with 29 and 28 PDZ binders [25,28] respectively, while about 50 PDZ proteins binds to the class I PBM of E6 viral oncoprotein from the human papillomavirus HPV16 [19]. By contrast, class II PBMs generally display lower affinities for their target and consequently a lowest number of binders in comparison with the class I PBMs. The hydrophobic character of the class II PBM sequences can explain the low affinities and sharp profiles of binding of SARS-CoV (6 binders) and SARS-CoV-2 (10 binders) E protein PBM with a reduced enthalpic contribution to the binding compared to the more hydrophilic class I sequences. Interestingly, the PBM of SARS-CoV-2 E protein can accommodate different classes of PDZ domains as illustrated in Table S2. In the case of the class III PBM of E protein for MERS-CoV, the large number of targets (29) can be mainly explained by the presence of a tryptophan residue at position - 1, a residue known to have a strong contribution to the binding affinity [29], potentially at the expense of the specificity. We have previously proposed that the large number of cell proteins targeted by WNV identified by Hold-up assay, could be related to the broad host repertoire of flaviviruses replicating in very different cells and species necessary for its complex zoonotic transmission cycle. By contrast, SARS-CoV viruses infect very specialized cells, such as bronchial epithelial cells, pneumocytes and upper respiratory tract cells in humans [30]. This observation can at least partially explain the narrow number of host proteins specifically targeted by SARS-CoV-2 during infection.

Six of the seven host PDZ partners of SARS-CoV E protein are also binders of SARS-CoV-2 E protein, in agreement with the strict conservation of their PBMs. However, the differences of sequence upstream the PBM enlarge the number of binders for SARS-CoV-2 protein E with four more host partners (Fig. 1A). Our results suggest that the SARS-CoV/SARS-CoV-2 PBM upstream sequence difference is responsible of the different binding profiles against the human PDZome. Several few studies indicate that residues up to the upstream eleventh position (p-11) can also be implicated in PDZ binding by modulating the affinity [29,31]. Thus, we can hypothesize that the substitution of Arg (p-6 in SARS-CoV sequence) by Glu (p-7 in SARS-CoV-2 sequence) may affect the potential electrostatic interaction of E protein with the ß2-ß3 loop of some PDZ domains. Interestingly, it has been reported that the substitution of a Glu at position - 5 with Arg in the inward rectifier K(+) channel protein GIRK3 disrupts its interaction with the PDZ domain of SNX27 [32]. Here, we also observed that SNX27 binds to SARS-CoV-2 E protein exposing an Arg at p-7, while SARS-CoV E protein coding for Glu at p-6 does not. This example illustrates the fine tuning of the affinity/specificity profiles by the four residues of the PBM, and the upstream sequence. Therefore, the charged residue substitution between SARS-CoV and SARS-CoV-2 PBM can modify the selective association of host PDZ proteins with the viral protein. The four host PDZ proteins, SNX27, RADIL, NHERF1, and MAST2 recognized by SARS-CoV-2, and not by SARS-CoV, can play a role in the differences observed between the two infections. The role of these PDZ proteins in viral pathogenesis remains to be explored.

#### Functions of host partners of SARS E protein

Among the five host proteins already reported as protein E partners (Schoeman et al., 2019), syntenin/SCDP and MPP5/PALS1 encode for a PBM potentially involved in PDZ binding. We did not detect the interaction between syntenin PDZ domain and SARS-CoV-2 E protein. With the holdup assay, we measured a weak interaction between MPP5/PALS1 PDZ domain and SARS-CoV-2 E PBM (Kd of 102 μM), while we did no monitored interaction between MPP5/PALS1 PDZ domain and SARS-CoV E PBM. Using equilibrium binding titration *in vitro* by Förster Resonance Energy Transfer (FRET), Toto and collaborators [33] have also calculated a better affinity of SARS-CoV-2 for MPP5/PALS1, illustrating the potential role of the upstream substitutions observed in the SARS-CoV-2 C-terminal E protein sequence in comparison with the SARS-CoV one. Teoh and collaborators [18] have proposed that this interaction between SARS-CoV E protein and MPP5/PALS1 alters the tight junctions in the lungs, MPP5/PALS1 being a key component of the Crumbs complex that controls the apical-basal polarity and tight junction formation. We cannot exclude that the rest of MPP5 and E proteins is required to stabilize their complex, as well as for syntenin that is reported to bind the E protein by strategies using full-length proteins (yeast two-hybrid, reciprocal coimmunoprecipitation and confocal microscopy assays).

Remarkably, we pinpointed that most of binding proteins are involved in cellular junctions and polarity, cellular trafficking and signaling. Indeed, our results highlighted that SARS-CoV-2 targets several compounds of the apical complex and tight/adherens junctions. Consequently, we expect that the SARS-CoV-2 proteins E, 3a and N can contribute to globally affect the cellular junctions and cellular polarity. We identified ZO-1 as the best binder for E and 3a proteins. ZO proteins provide a network of scaffold proteins that tether the transmembrane tight junction proteins, comprising the Claudins, to the actin cytoskeleton [34]. The three ZO paralogous proteins, ZO-1, ZO-2 and ZO-3, encode for three PDZ domains in their N-terminal region. In our *in vitro* assay, the second PDZ of ZO-1/TJP1 displays the highest affinity for E and 3a proteins of SARS-CoV2 in the micromolar range. This domain has been previously described as the binding site of ZO-1 for the connexin43 [35], involved in the organization of the gap junction, but also in the ZO-2 and ZO-3 heteroassociation. Dimerization of ZO-1 occurs through the interaction between ZO1-PDZ2 domains and the swapped-dimerization of ZO-1-PDZ2 regulates the connexin-43 binding [36]. Interestingly, ZO proteins also interact with Afadin/MLLT4 while the PBM sequence of ZO-2 binds to the PDZ domain of SNX27 to regulate the trafficking of ZO-2 to the tight junction [37,38]. PARD3 is also involved in the formation of adherens and tight junctions. Here, we showed that MLLT4, SNX27, PARD3, and MPP5/PALS1 to a lesser extent, are also targeted by SARS-CoV-2 E PBM. By interacting by at least five proteins directly involved in the organization of the tight junction complexes, we expect a global perturbation of the epithelial/endothelial barriers by SARS-CoV-2 during infection. In this direction, SARS-CoV-2 replication causes a transient decrease in cellular barrier function and disruption of tight junctions [39] and a SARS-CoV-2 productive infection locally alters the distribution of ZO-1, which associates to tight junctions. The characteristic ZO-1 staining pattern at cell boundaries appeared disrupted in areas with viral expression, suggesting a possible impairment of epithelial barrier integrity.

Targeting host proteins from polarity and junctions complexes may have a consequence on the morphology of epithelium/endothelium but also on the immune response and cellular homeostasis [40]. In addition to their structural role in many tissues and organs, they can also regulate signaling mechanisms. Tight junctions are linked to many human diseases. Therefore, chronic inflammatory diseases and cancer have been related to dysfunction of tight junctions. MPP5/PALS1 of the Crumbs complex is expressed in T lymphocytes and is required for the T Cell-receptor (TCR)-mediated activation [41]. ZO-1 and ZO-2 are also expressed in T lymphocytes and are relocated to the immune synapse after TCR stimulation. SNX27 regulates trafficking and recycling in T lymphocytes and controls ZO-2 mobility at the immune synapse through a PDZ-dependent interaction [42]. LNX1/2 is involved in signaling with involvement in ubiquitination subsequent proteasomal degradation. The PDZ3 domain of PARD3 binds to the transcription factor YAP (Yes-associated protein) [43] regulating the activation of the Hippo pathway, a signaling pathway modulating cell proliferation and cell death. YAP protein can be found associated to the adherens and tight junctions and several regulators of Hippo pathway are components of the the adherens and tight junctions [44]. Finally, NHERF1 regulates the Wnt/b-catenin signaling, a pathway that regulates cell growth and proliferation. NHERF1 has been shown to be targeted for degradation by E6 from HPV [45]. RHPN2 (Rho pathway; [46]), MAST2 (anti-survival pathway, PTEN/Akt signaling; [12]), RGS3 (G protein signaling; [47]), PTPN13 (tumor suppressor, Akt signaling; [48]) and HTRA1 (Signaling pathway TGF-ß; [49]) play also function in signaling pathways. All these proteins have been found to significantly interact with SARS-CoV-2 proteins in our hold-up assay. Seven of the 14 host targets of SARS-CoV2 proteins E, 3a, and N significantly impacted viral replication. We showed that knockdown of the PDZ-containing proteins PARD3 and RSG3 results in a decrease of SARS-CoV-2 replication in cells while the silencing of MLLT4, PARD3B, MAST2, RHPN2, and HTRA1 increase virus replication. These proteins are involved in the regulation of cellular junction and polarity, but also in cellular signaling. However, we cannot conclude on the PDZ/PBM dependence of this result and further in cell studies are needed to decipher the functional role of these proteins in the viral cycle of SARS-CoV-2.

#### PDZ proteins recognized by SARS-CoV-2 E protein are also host targets of others viruses

Many studies provide structural and biological evidences that the viral proteins act as competitors endowed with specificity and affinity in an essential cellular process by mimicking PBM of cellular partners. Disruption of critical endogenous protein–protein interactions by viral protein drastically alters intracellular protein trafficking and catalytic activity of cellular proteins that control cell homeostasis. Importantly, most of our selected host PDZ-containing proteins have been previously reported as host proteins targeted by viruses. Tight junctions are known to be targeted by multiple pathogenic viruses [50,51]. PARD3 and TJP1 are targets of flavivirus proteins (WNV, TBEV and DV) [28,52–54]. Others cellular PDZ proteins were also identified as targets for other virus including DLG1 for hepatitis C virus and influenza virus [55,56]; NHERF1, HTRA1, PARD3, SNX27 for human papillomavirus [45,57–59].

Here, we showed that the three viral PBM-containing proteins expressed by SARS-CoV-2 target several cellular proteins involved in the maintenance and regulation of the tight junctions and immune response. By hijacking junctional proteins and signaling mechanisms, viruses ensure their propagation through the manipulation of the cellular homeostasis, the immune system and tissue barriers. The perturbation of cellular fitness promotes their replication and spread. Many PDZ-containing proteins are involved in cell-cell junctions, cell polarity, recycling, or trafficking. Mimicking short linear PBM motifs to target PDZ protein and subvert host functionalit*y* is a high adaptation capacity and a significant evolutionary advantage of many viruses [40]. We previously showed that the PBM from the glycoprotein encoded by the rabies virus competes with PTEN for binding to the anti-survival kinase MAST2 and drastically affects the PTEN/Akt signaling pathway. Consequently, the viral glycoprotein/MAST2 interaction promotes survival of the infected neurons to ensure viral propagation. As already observed for other viruses, potential pleiotropic perturbation of cellular functions can be induced by SARS-CoV-2 during infection comprising cell survival/death (MAST2, PARD3, NHERF1), immune response (MPP5/PALS1, syntenin), and cellular organization (ZO-1, MLLT4).

## Acknowledgements

This work was supported by the URGENCE COVID-19 fundraising campaign of Institut Pasteur, the ANR Recherche Action Covid19–FRM PDZCov2 program and by the French Infrastructure for Integrated Structural Biology (FRISBI) ANR-10-INSB-05-01 for AFMB. G. G. is a recipient of the Post-doctorants en France program of the Fondation ARC fellowship.

The authors acknowledge Gilles Travé for helpful discussion and for proofreading of the manuscript.

## Materials and methods

### Peptide synthesis

The peptides were synthesized in solid phase using Fmoc strategy (Proteogenix) encompassing a biotinyl group, a (PEG)3 or 6-Aminohexanoic acid (Ahx) spacer and the C-terminal PBM sequence of E, 3A and N proteins (12 residues long). Peptides were resuspended in water or in a mix of DMSO and buffer and used for holdup and competitive fluorescence polarization assays.

### Holdup assays

We used an updated version of our original PDZome library [19], that contains 266 soluble PDZ [20]. This PDZome v2 library was prepared as previously described [20].

The holdup assay was carried out with the biotinylated viral PBMs against the full human PDZ library in two independent experiments[19,20]. Error bars are standard deviations of the two experiments. Affinity values (Figure 1, Table 1) were calculated from the Binding Intensities according to the method published in Gogl et al. [23]. Immobilized peptide concentrations were calculated using measured dissociation constants from competitive fluorescent polarization assays (see below). We have found that all determined BI-Kd pairs resulted a mean peptide concentration of ~20 μM, a concentration that is coherent with other peptides in the same method [22,23,60]. The minimal Binding Intensity (BI) threshold value is 0.2 to define a significant interaction, roughly corresponding to a 100 μM dissociation constant as previously reported [19].

### Competitive fluorescence polarization assay

Competitive fluorescence polarization measurement and data evaluation were performed as described by Simon and collaborators [61]. Four different fluorescent peptides were used f16E6 (fluorescein-RTRRETQL), fRSK1 (fluorescein-KLPSTTL), fpRSK1 (fluorescein-KLPpSTTL), and MERS-CoV. The MERS-CoV peptide was labeled with sub-stoichiometric FITC (Sigma-Aldrich, St. Louis, Missouri) in a basic HEPES buffer (pH 8.2) and the reaction was stopped with 100 mM TRIS. The peptide was buffer exchanged in order to remove fluorescent contaminants. Fluorescence polarization was measured with a PHERAstar (BMG Labtech, Offenburg, Germany) microplate reader by using 485 ± 20 nm and 528 ± 20 nm band-pass filters (for excitation and emission, respectively). In direct FP measurements, a dilution series of the Ni- and Amylose-affinity purified MBP-PDZ proteins was prepared in a 20 mM HEPES pH 7.5 buffer that contained 150 mM NaCl, 0.5 mM TCEP, 0,01% Tween 20 and 50 nM fluorescently labeled peptide. In total, the polarization of the probe was measured at 8 different protein concentrations. In competitive FP measurements, the same buffer was supplemented with the protein to achieve a complex formation of 60-80%, based on the direct titration. Then, this mixture was used for creating a dilution series of the competitor (i.e. the studied peptides) and the measurement was carried out identically as in the direct experiment. Analysis of FP experiments were carried out using ProFit.

### Cells

A549 cells stably expressing ACE2 (A549-ACE2, kindly provided by Dr. Olivier Schwartz), were propagated at 37°C, 5% CO2 in DMEM with L-glutamine (Gibco) supplemented with 10% FBS, penicillin-streptomycin and 20 μg/mL blasticidin S.

### Virus

The SARS-CoV-2 strain used (BetaCoV/France/IDF0372/2020 strain) is a kind gift from the National Reference Centre for Respiratory Viruses at Institut Pasteur, Paris, originally supplied through the European Virus Archive goes Global platform. It was propagated once in Vero-E6 cells.

### siRNA

#### transfections

An OnTargetPlus siRNA SMARTpool library (Horizon Discovery) was purchased targeting 20 human PDZ-containing proteins (Table S3). This library was arrayed in 96-well format, and also included non-targeting siRNAs and siRNA pools targeting ACE2. The siRNA library was transfected into A549-ACE2 cells seeded in a 384-well plate format (6250 cells per well), using Lipofectamine RNAiMAX reagent (Thermo Fisher).

Briefly, 4 pmoles of each siRNA pool were mixed with 0.1 μL RNAiMAX transfection reagent and OptiMEM (Thermo Fisher) in a total volume of 10 μL. After a 5-minute incubation period, the transfection mix was added to cells. 48 h post transfection, the cells were either subjected to SARS-CoV-2 infection or incubated for 72 hours to assess cell viability using the CellTiter-Glo luminescent viability assay, following the manufacturer’s instructions (Promega). Luminescence was measured in a Tecan Infinity 2000 plate reader, and percentage viability calculated relative to untreated cells (100% viability) and cells lysed with 20% ethanol or 4% formalin (0% viability), included in each experiment.

### Virus infections

For infections, the cell culture medium was replaced with virus inoculum (MOI 0.1 PFU/cell) and incubated for one hour at 37°C, 5% CO2. After one hour adsorption period, the inoculum was removed, replaced with fresh media, and cells incubated at 37°C, 5% CO2. 72 h post infection, the cell culture supernatant was harvested, and viral load assessed by RT-qPCR. Briefly, the cell culture supernatant was collected, heat inactivated at 95°C for 5 minutes and used for RT-qPCR analysis. SARS-CoV-2 specific primers targeting the N gene region: 5’-TAATCAGACAAGGAACTGATTA-3’ (Forward) and 5’-CGAAGGTGTGACTTCCATG-3’ (Reverse) were used with the Luna Universal One-Step RT-qPCR Kit (New England Biolabs) in an Applied Biosystems QuantStudio 6 thermocycler, with the following cycling conditions: 55°C for 10 min, 95°C for 1 minute, and 40 cycles of 95°C for 10 seconds, followed by 60°C for 1 minute. The number of viral genomes is expressed as PFU equivalents/mL and was calculated by performing a standard curve with RNA derived from a viral stock with a known viral titer. The effect of knock-downs on virus replication was compared to control siRNA-transfected cells in three independent experiments with Dunnett’s multiple comparison test based on a two-way ANOVA.

