## Supplemental Tables S1, S2 & S3 for "Host PDZ-containing proteins targeted by SARS-Cov-2"

**Table S1;** All BI values obtained by holdup assay using PBM peptides from SARS-CoV, SARS-Cov-2, and MERS-CoV E proteins, and from SARS-CoV-2 N and 3a proteins. Kd values are calculated for BI values higher than 0.1.

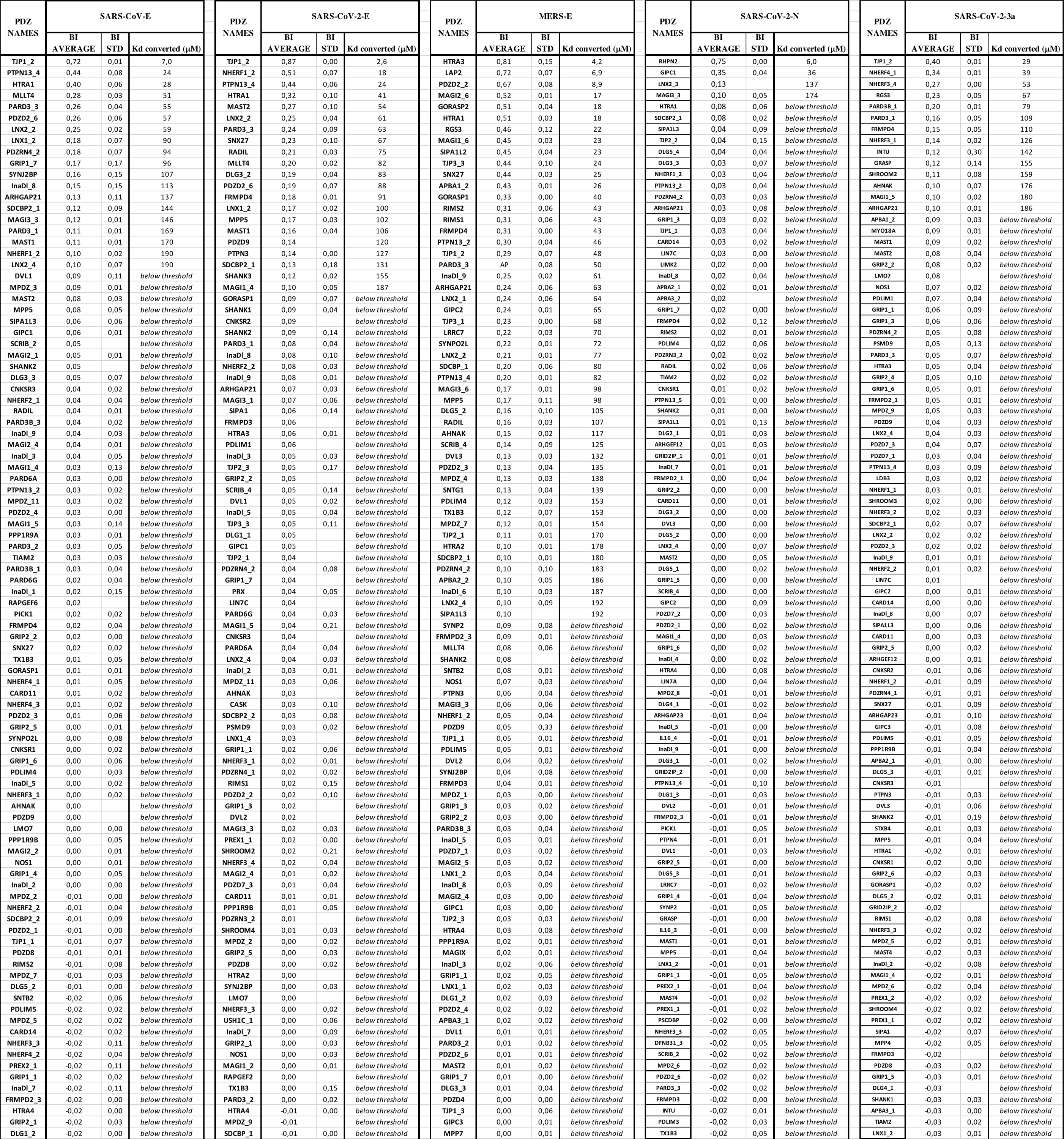

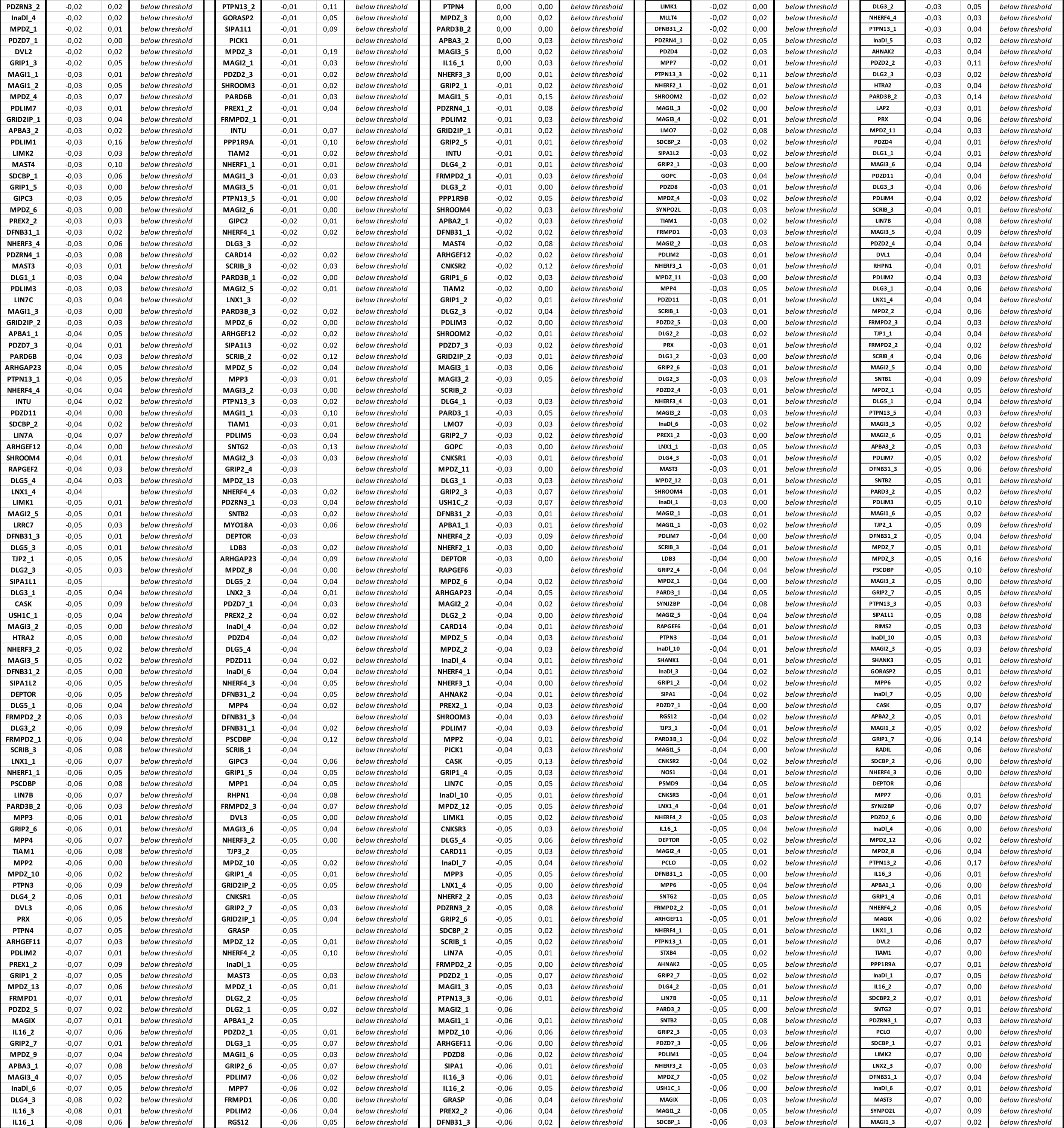

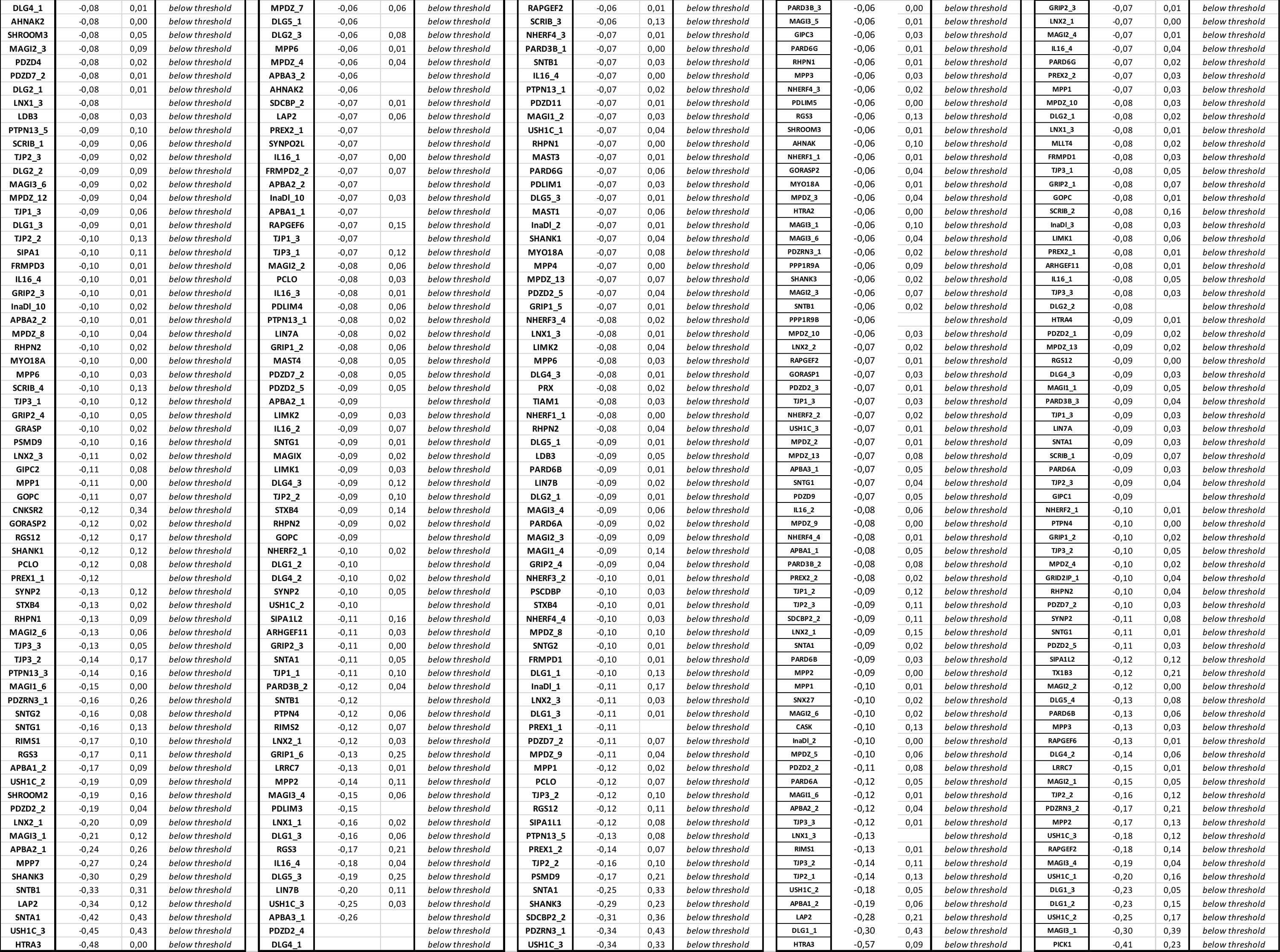

**Table S2; Classes of PDZ Domains targeted by the SARS-CoV-2 E protein PBM.**

**Left panel;** name of selected PDZ domain, PDB code of their structure and the residue of PDZ helix α2 in interaction with p-2 of PBM.

**Right panel**; structural view of the PBM binding groove with the residue of helix a2 facing the position -2 of PBM highlighted in blue sticks.

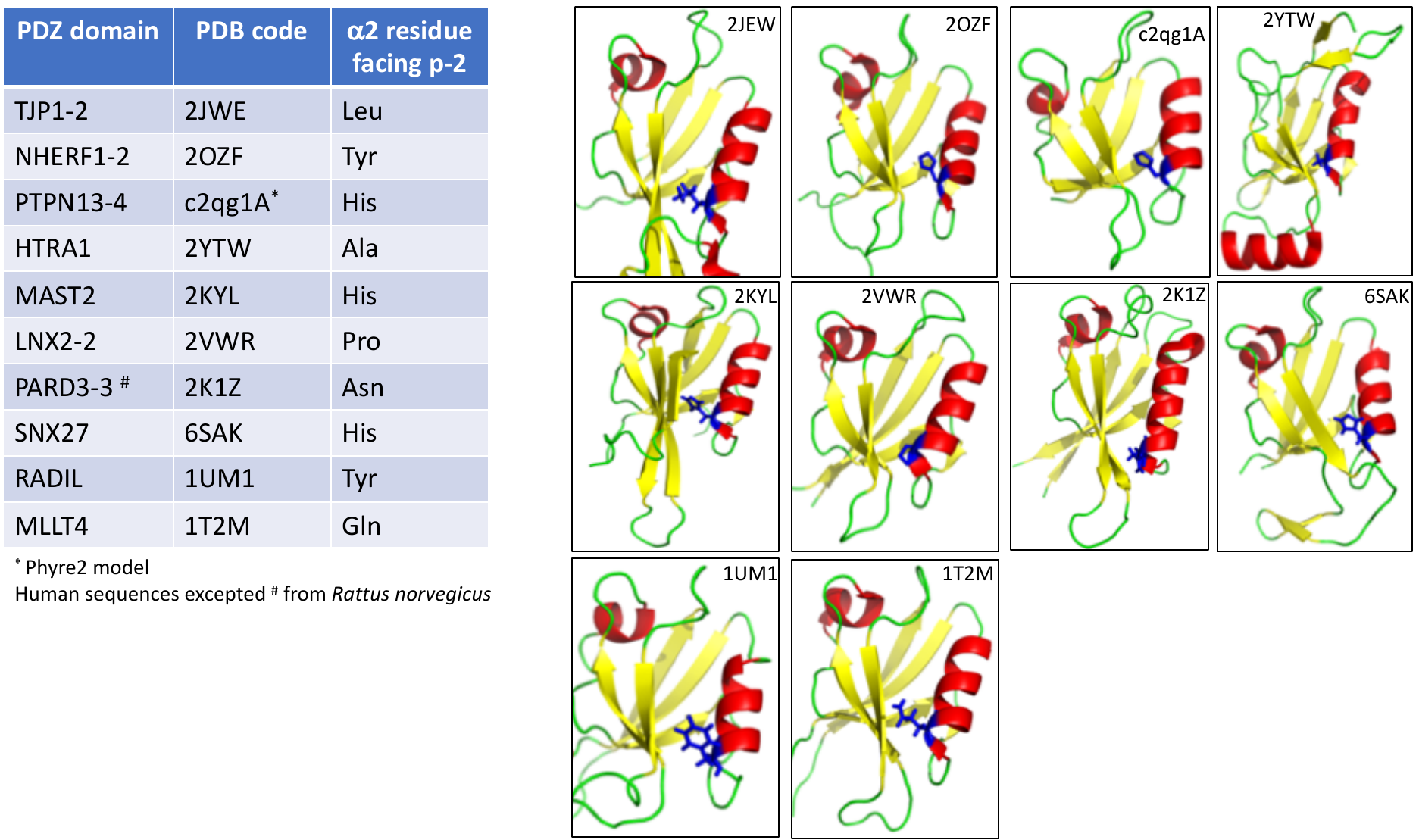

**Table S3;** OnTargetPlus siRNA SMARTpool library (Horizon Discovery) was purchased targeting the 20 selected human PDZ-containing proteins.

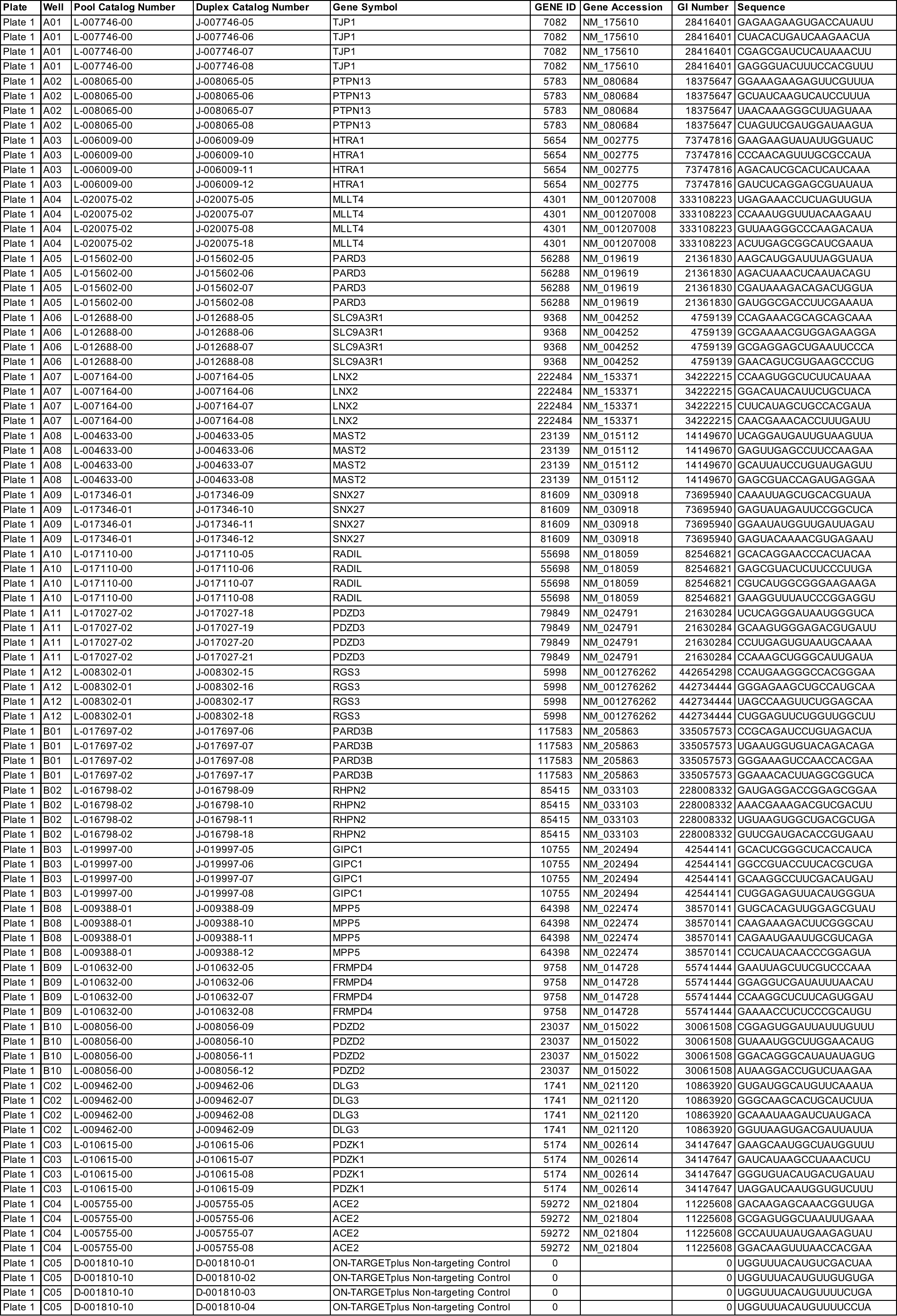
